# SVIM-asm: Structural variant detection from haploid and diploid genome assemblies

**DOI:** 10.1101/2020.10.27.356907

**Authors:** David Heller, Martin Vingron

**Affiliations:** Computational Molecular Biology Department, Max Planck Institute for Molecular Genetics, Berlin, 14195, Germany

## Abstract

**Motivation:** With the availability of new sequencing technologies, the generation of haplotype-resolved genome assemblies up to chromosome scale has become feasible. These assemblies capture the complete genetic information of both parental haplotypes, increase structural variant (SV) calling sensitivity and enable direct genotyping and phasing of SVs. Yet, existing SV callers are designed for haploid genome assemblies only, do not support genotyping or detect only a limited set of SV classes.

**Results:** We introduce our method SVIM-asm for the detection and genotyping of six common classes of SVs from haploid and diploid genome assemblies. Compared against the only other existing SV caller for diploid assemblies, DipCall, SVIM-asm detects more SV classes and reached higher F1 scores for the detection of insertions and deletions on two recently published assemblies of the HG002 individual.

**Availability and Implementation:** SVIM-asm has been implemented in Python and can be easily installed via bioconda. Its source code is available at github.com/eldariont/svim-asm.

**Supplementary information:** Supplementary data are available online.

## 1 Introduction

As one of the main classes of genomic variation, structural variants (SVs) comprise a diverse range of genomic rearrangements with sizes larger than 50 bps. Although there are considerably less SVs than Single Nucleotide Variants (SNVs) or small indels in an average human genome, SVs affect more base-pairs (1000 Genomes Project Consortium, 2015). Consequently, SVs have a strong effect both on the healthy human phenotype and human disease.

Due to the availability of affordable and accurate next-generation sequencing (NGS) technology, SVs are now commonly detected by the analysis of sequencing reads. Typically, the reads from a genome under investigation are aligned to an existing reference genome to reveal differences between both genomes (read-based SV calling). Alternatively, de novo assembly uses sequence overlaps between reads to computationally reconstruct longer genomic fragments, called contigs. Like raw sequencing reads, these assembly contigs can be aligned to a reference or comparison genome to facilitate the detection of SVs (assembly-based SV calling) (Sedlazeck et al., 2018).

The detection of SVs from contigs instead of raw reads is particularly valuable for the analysis of species for which no high-quality reference genome is available. Other applications of assembly-based SV calling include the analysis of genomes with large-scale rearrangements compared to the reference, the pair-wise comparison of multiple related genome assemblies and the analysis of sample-specific sequences or large insertions. While a growing number of software tools detect SVs from the alignments of short and long reads (Kosugi et al., 2019), only few tools are available for the detection of SVs from genome-genome alignments. Three such tools, AsmVar, Assemblytics and SyRy, have been developed recently but only for the analysis of haploid genome assemblies (Goel et al., 2019).

Until recently, genome assembly methods usually collapsed the two parental haplotypes of a diploid genome into a haploid genome representation. With the availability of longer sequencing reads and complementary sequencing technologies like Hi-C and Strand-Seq, however, the routine production of haplotype-resolved genome assemblies has become feasible (Garg et al., 2019; Nurk et al., 2020). These haplotype-resolved assemblies capture the complete genetic information of both parental haplotypes, can increase variant calling sensitivity (Chaisson et al., 2019) and enable direct genotyping and phasing of variants. Yet, only one method, DipCall, has been published so far for the detection of large insertions and deletions from haplotype-resolved genome assemblies (Li et al., 2018). In this study, we introduce our method SVIM-asm for the detection and genotyping of six common classes of SVs, including insertions and deletions, from haploid and diploid genome assemblies.

## 2 Materials and Methods

SVIM-asm (Structural Variant Identification Method for Assemblies) is based on our previous method SVIM that detects SVs in long-read alignments (Heller and Vingron, 2019). Although SVIM-asm follows a similar workflow as SVIM, several adaptions have been made to consider the unique properties of assembly alignments compared to read alignments (see Figure S1). To enable the analysis of both haploid and diploid genome assemblies, SVIM-asm implements two alternative pipelines (see Figure S2).

Diploid genome assemblies consist of two sets of contigs, one for each parental haplotype. In the first step of the pipeline (*COLLECT*), SV signatures are extracted separately for each haplotype from discordant alignments of individual contigs to the reference. The discordancies fall into two categories: a) long alignment gaps within alignment segments (intra-alignment discordancies) and b) discordant positions and orientations between alignment segments (inter-alignment discordancies).

In the second step of the pipeline (*PAIR*), signatures from opposite haplotypes are compared and paired up if sufficiently similar. To measure the similarity of two signatures, the edit distance (Levenshtein distance) between their haplotype sequences is computed with the library edlib (Šošić and Šikić, 2017). Based on the computed distances, very similar signatures (i.e. signatures with similar haplotype sequences) from different haplotypes are merged.

In the third step of the pipeline (*GENOTYPE*), paired signatures from the two opposite haplotypes are merged into homozygous SV candidates while variants without a partner on the other haplotype are called as heterozygous SV candidates. Finally, the genotyped SVs are written out in Variant Call Format (VCF) as members of one of six SV classes (*OUTPUT*).

In contrast to their diploid counterparts, haploid assemblies consist of only a single set of contigs. For diploid organisms, this set often represents a mixture of the two haplotypes. Due to the missing second haplotype, it is not possible to estimate genotypes from haploid genome assemblies. After the same first step (*COLLECT*) is applied to the assembly alignments, the *PAIR* and *GENOTYPE* steps are skipped for haploid assemblies and the detected SV signatures are written out immediately (*OUTPUT*).

## 3 Results

We compared our tool, SVIM-asm (v1.0.0), to the DipCall pipeline (v0.1). For the evaluation we chose two publicly available diploid genome assemblies of the HG002 individual from Wenger et al. (Assembly A) and Garg et al. (Assembly B) (see Supplementary Methods) (Wenger et al., 2019; Garg et al., 2019). We aligned the fragments separately for each haplotype using minimap2 (v2.17-r941) (Li, 2018) and produced genotyped SV calls using SVIM-asm and DipCall, respectively.

When compared against the GIAB SV benchmark set of 7,281 insertions and 5,464 deletions (Zook et al., 2020) using truvari (v2.0.1), both methods reached F1 scores above 90% (see Figure S3, upper panel). SVIM-asm performed slightly better than DipCall with F1 scores of 93.2% (Assembly A) and 93.7% (Assembly B) compared to 91.7% and 92.5%, respectively. This improvement was enabled by a smaller number of false positives (violet) and false negatives (blue) in the SVIM-asm callset (see Figure 1). When measuring precision and recall across variant lengths, we observed that SVIM-asm reached a higher recall than DipCall particularly for large deletions and insertions (see Figure S4). We attribute this to inter-alignment discordancies from split alignments at large variants which are analyzed by SVIM-asm but ignored by DipCall.

**Figure 1:**
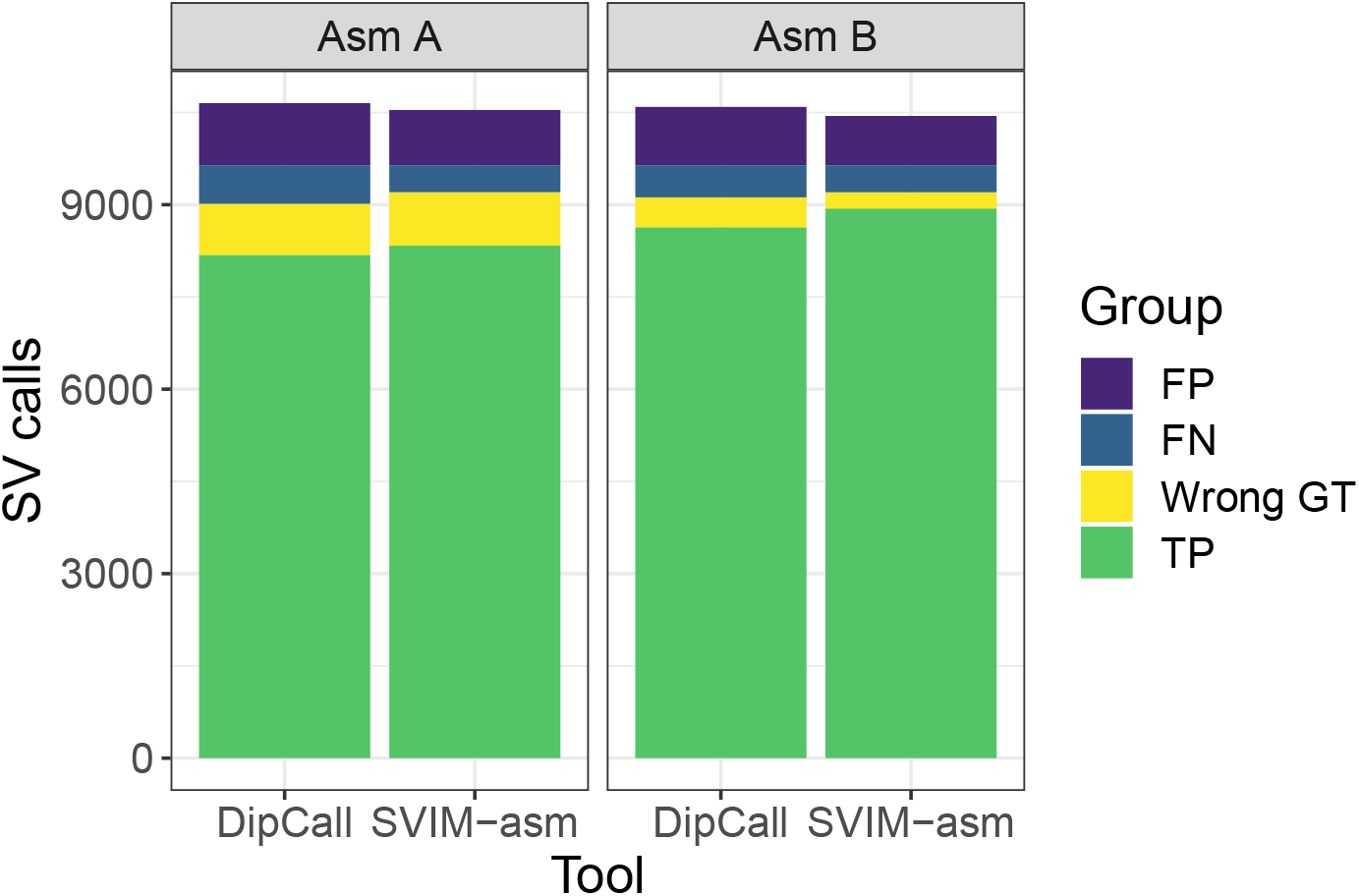
Comparison of SV detection and genotyping performance of DipCall and SVIM-asm (x-axis) on two diploid genome assemblies (left panel: Asm A by Wenger et al. and right panel: Asm B by Garg et al.). This stacked barplot shows the number of true positives with correct genotype (green), true positives with wrong genotype (yellow), false negatives (blue) and false positives (violet) as defined by comparing against the GIAB SV benchmark set. For a plot of precision, recall and F1-score, see Figure S3.

We also analyzed the genotypes of true positive calls and in general observed a very high concordance above 90% with the benchmark set. For both assemblies, SVIM-asm reached a higher number of true positives with correct genotype (green, Figure 1) than DipCall. When requiring true positives to have correct genotypes, SVIM-asm reached F1 scores of 84.4% (Assembly A) and 91.0% (Assembly B) while DipCall only reached 83.2% and 87.6%, respectively (see Figure S3, lower panel).

Compared to DipCall which detects only insertions and deletions, SVIM-asm additionally calls tandem and interspersed duplications, inversions and translocation breakends. As defined in the VCF for the specification of complex rearrangements, each translocation breakend represents one side of a novel adjacency between two distant genomic loci. From the Assemblies A and B SVIM-asm detected 14,399 / 13,929 insertions, 9,407 / 9,154 deletions, 89 / 72 inversions, 109 / 99 tandem duplications, 2 / 4 interspersed duplications, and 376 / 1,340 translocation breakends, respectively (see Figures S5 and S6).

## 4 Discussion

The detection of structural variants from genome assemblies complements read-based SV calling approaches, allows the pairwise comparison of genomes and enables SV calling even in the absence of a suitable reference genome. In this study, we introduced SVIM-asm, an accurate software tool for the detection of SVs from haploid and diploid genome assemblies. Compared to existing tools for assembly-based SV detection, SVIM-asm supports more SV types, reached a higher SV calling performance in our benchmarks and predicted genotypes more precisely.

## Acknowledgements

The authors wish to thank Shilpa Garg for encouraging the application of SVIM to genome assemblies and for providing the diploid genome assembly dataset ahead of publication.

## Funding

This work was supported by the German Bundesministerium für Bildung und Forschung (031L0169A) and the doctoral program of the International Max Planck Research School for Biology and Computation.

## Notes

### Competing Interest Statement

The authors have declared no competing interest.

### Summary of Updates

Changed tone of introduction and discussion to avoid impression that SVIM-asm would be a replacement for read-based SV callers like pbsv; Updated evaluation results for genotyped SV calls; Transformed Figure 1 into stacked barplot of different error types; Updated Figures S5 and S6 to only show variants detected in chromosomal contigs

