## Supplementary Material for "SVIM-asm: Structural variant detection from haploid and diploid genome assemblies"

*David Heller and Martin Vingron*
*Computational Molecular Biology Department, Max Planck Institute for Molecular Genetics, Berlin, 14195, Germany*

### Supplementary Figures

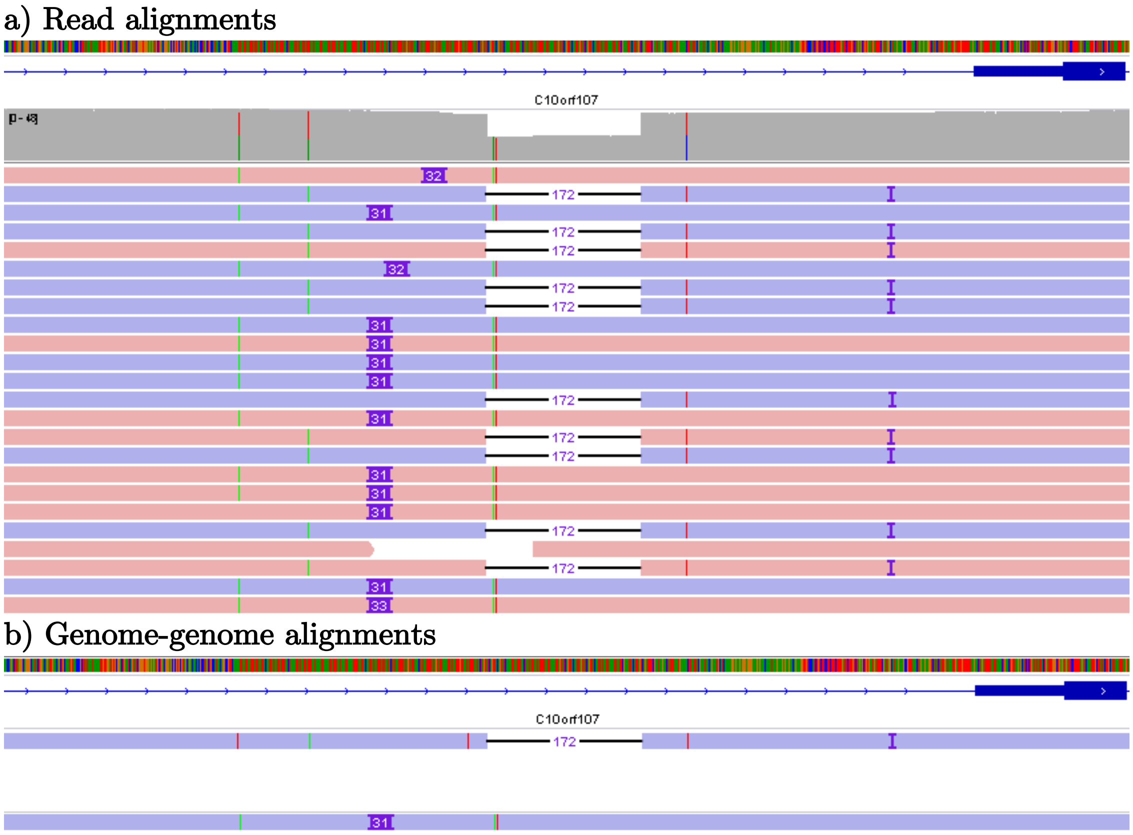


**Fig. S1 – Alignments of sequencing reads versus genome assemblies:** Both sequencing reads (panel a) and contigs from a genome assembly (panel b) can be aligned to a reference genome.

**a** | Several long reads are aligned to a reference genome on the forward (red) or reverse strand (blue). Approximately half of the reads belong to one parental haplotype containing a 172 bp deletion while the other half belong to the other parental haplotype containing an insertion of approximately 31 bp instead.

**b** | A diploid genome assembly consisting of two sets of contigs is aligned to the same genomic region. One contig (upper alignment) represents the haplotype carrying the deletion while the other contig (lower alignment) represents the haplotype with the insertion.

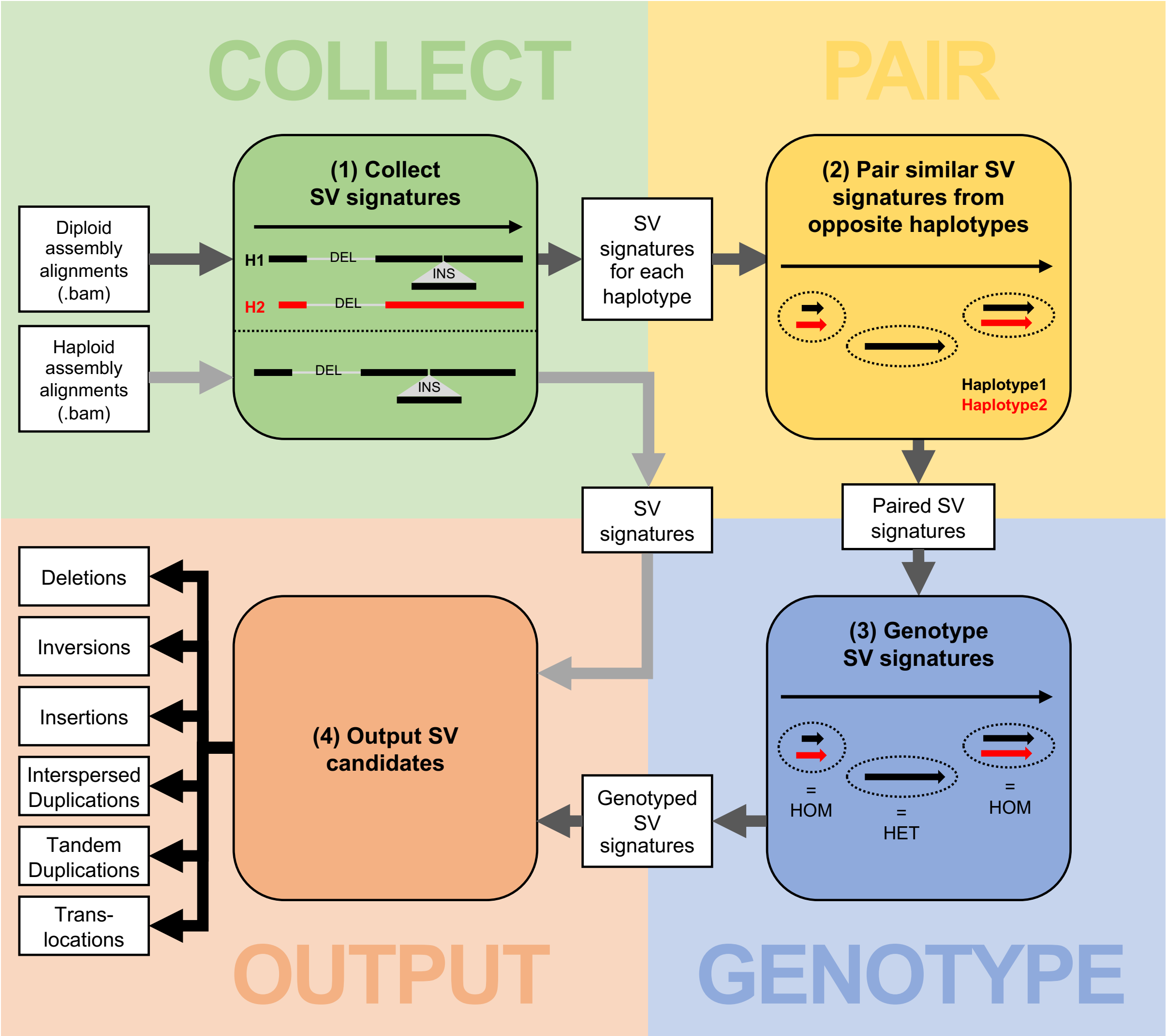


**Fig. S2 - The workflow of SVIM-asm:** Signatures for SVs are collected from the input assembly alignments (COLLECT). Depending on the ploidy of the assembly, signatures are collected from a single haplotype (haploid assembly) or a pair of haplotypes (diploid assembly). For diploid assemblies, SV signatures from the two haplotypes are compared and similar signatures are paired up based on the edit distance between their haplotype sequences (PAIR). Paired SV signatures represent homozygous (HOM) SVs while isolated SV signatures from only one of the haplotypes represent heterozygous (HET) SVs (GENOTYPE). Finally, six different classes of SV candidates are written out in VCF: deletions, insertions, inversions, interspersed duplications, tandem duplications and translocations (OUTPUT). SV signatures from haploid assemblies skip steps PAIR and GENOTYPE and are written out directly after the COLLECT step.

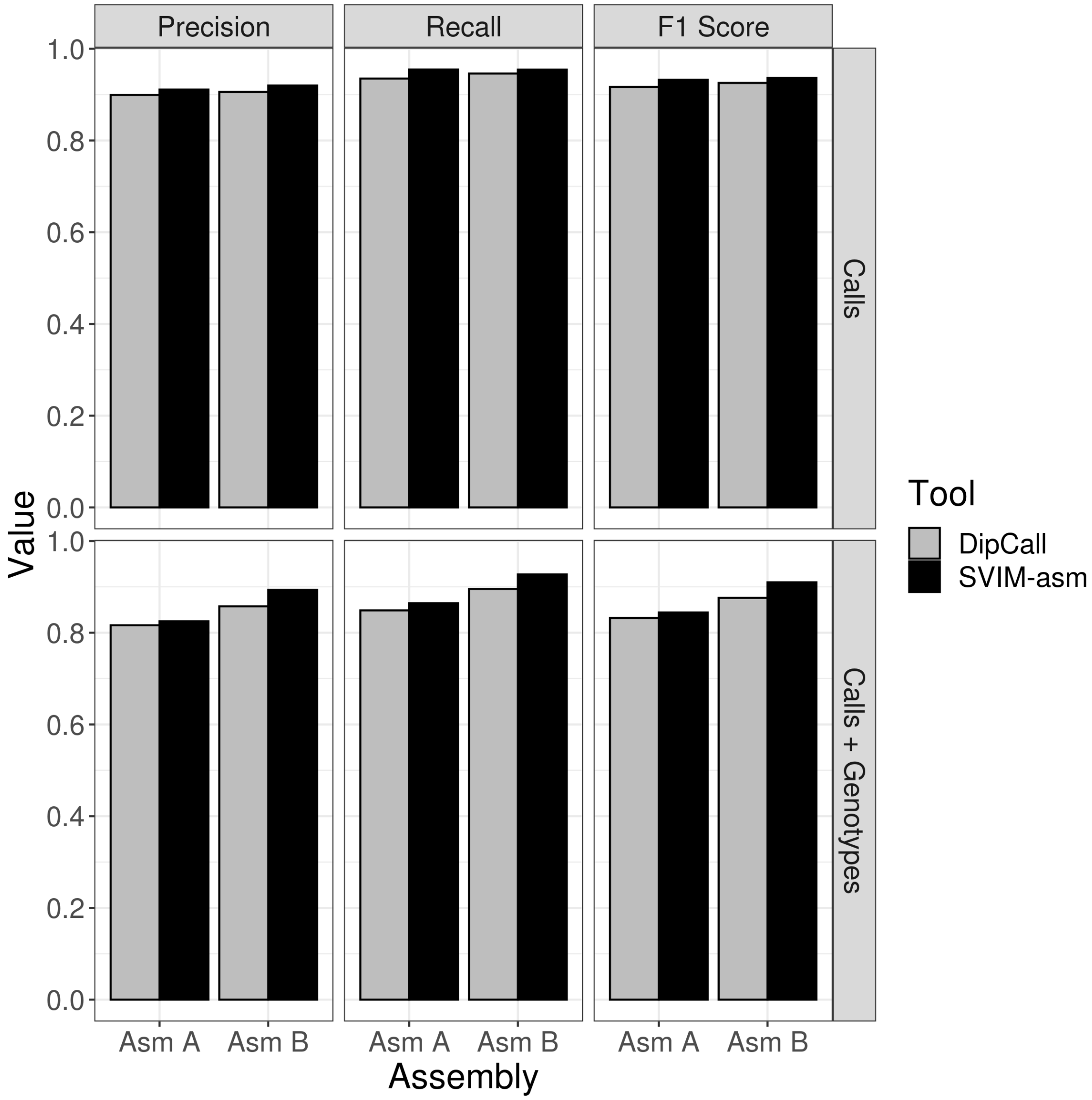


**Fig. S3 – Comparison of SV detection performance of DipCall and SVIM-asm.** For two diploid genome assemblies (x-axis, Asm A by Wenger et al. and Asm B by Garg et al.) the precision, recall and F1 score (y-axis) were measured by comparing against the GIAB SV benchmark set. In the uppel panel “Calls”, the ability of the tools to detect the presence of SVs regardless of their genotype was evaluated, i.e. SV calls were classified as true positives regardless of their genotype. In the lower panel “Calls + Genotypes”, only SV calls with identical genotypes in the benchmark set were classified as true positives.

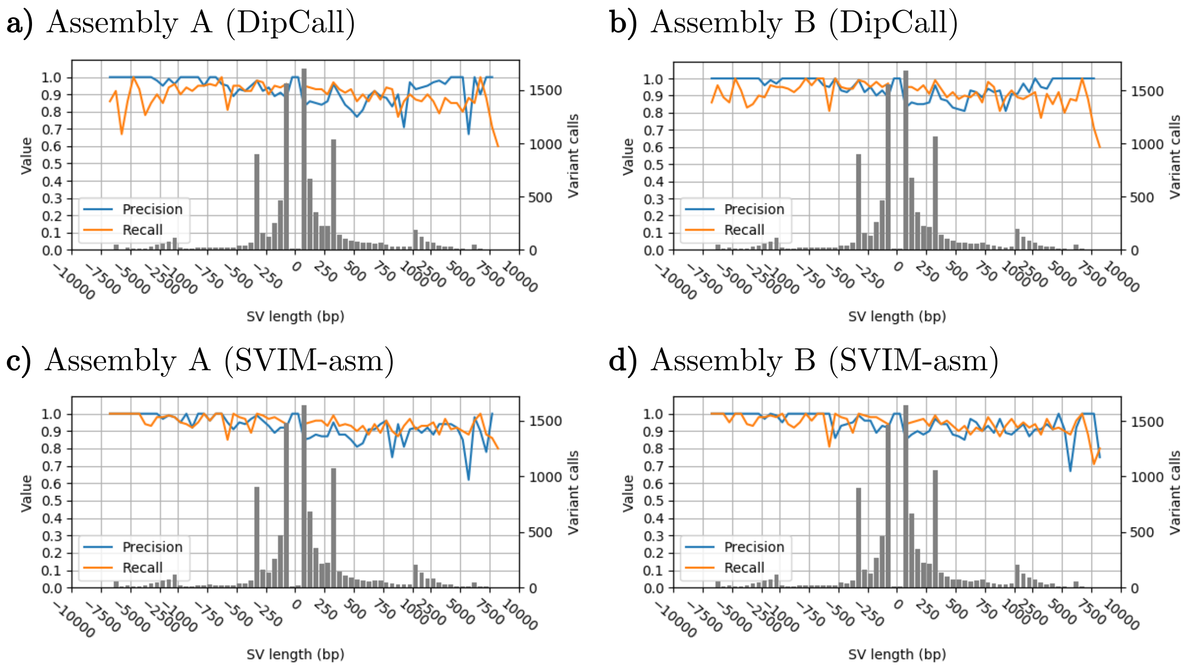


**Fig. S4 – SV detection performance across variant lengths:** Precision, recall and number of variant calls (y-axis) are shown for different SV length bins (x-axis) for the SV callers DipCall (a and b) and SVIM-asm (c and d). SV calls were classified as true positives regardless of their genotype. The bin size is 50 bp for variants shorter than 1 kbp, and 500 bp for variants >1 kbp. Positive lengths indicate insertions and negative lengths indicate deletions.

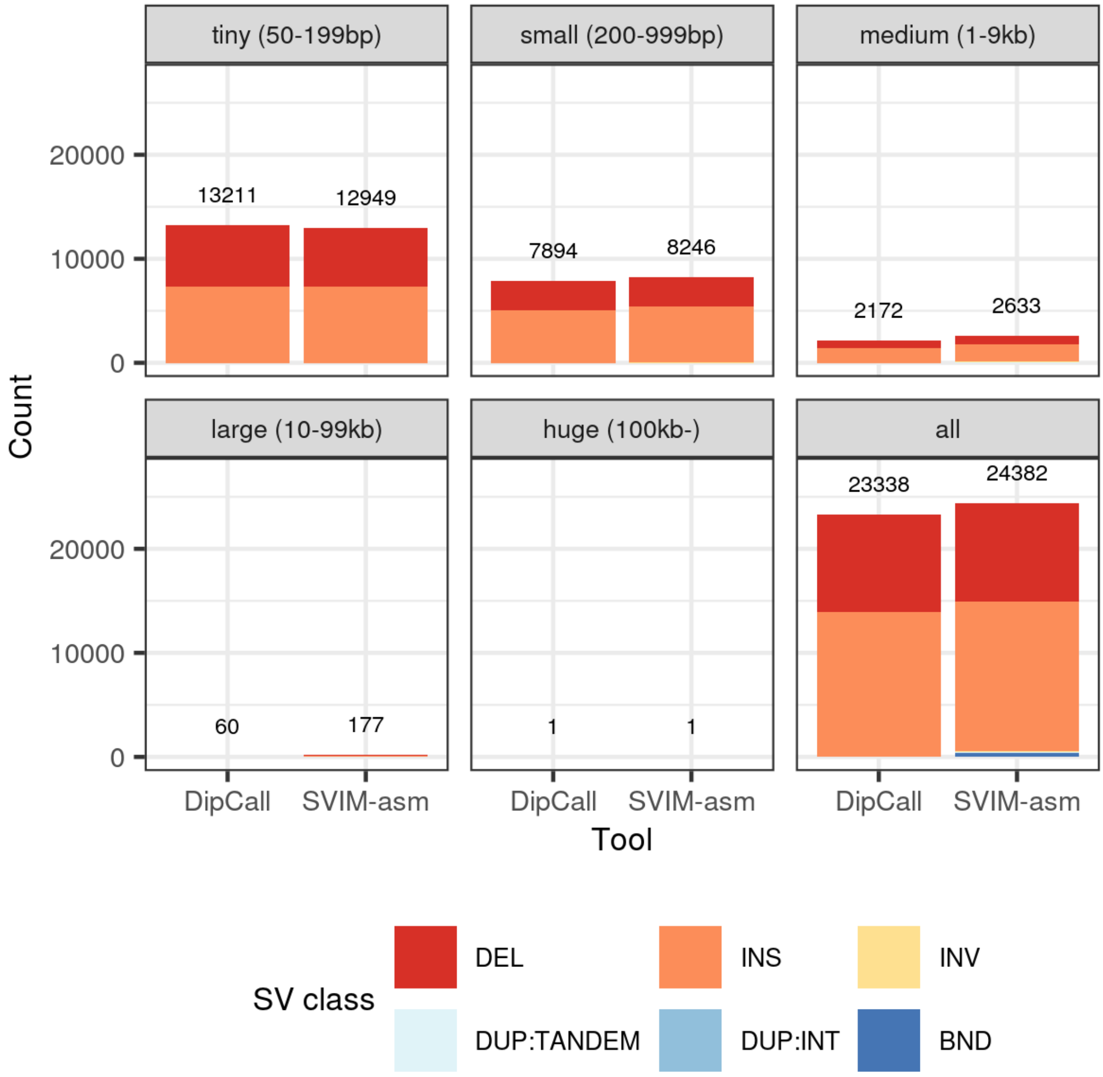


**Fig. S5 – Number of SV calls from Assembly A grouped into five size classes:** Shown are stacked bar plots of SV classes represented by different colors. Each panel represents one size class and visualizes the number of calls in that size range called by DipCall and SVIM-asm within the chromosomal contigs (1 through 22 plus X, Y and MT). The bottom right panel shows the counts for all SV calls regardless of size. As translocation breakends (BND) do not have a size, they are included only in this bottom right panel.

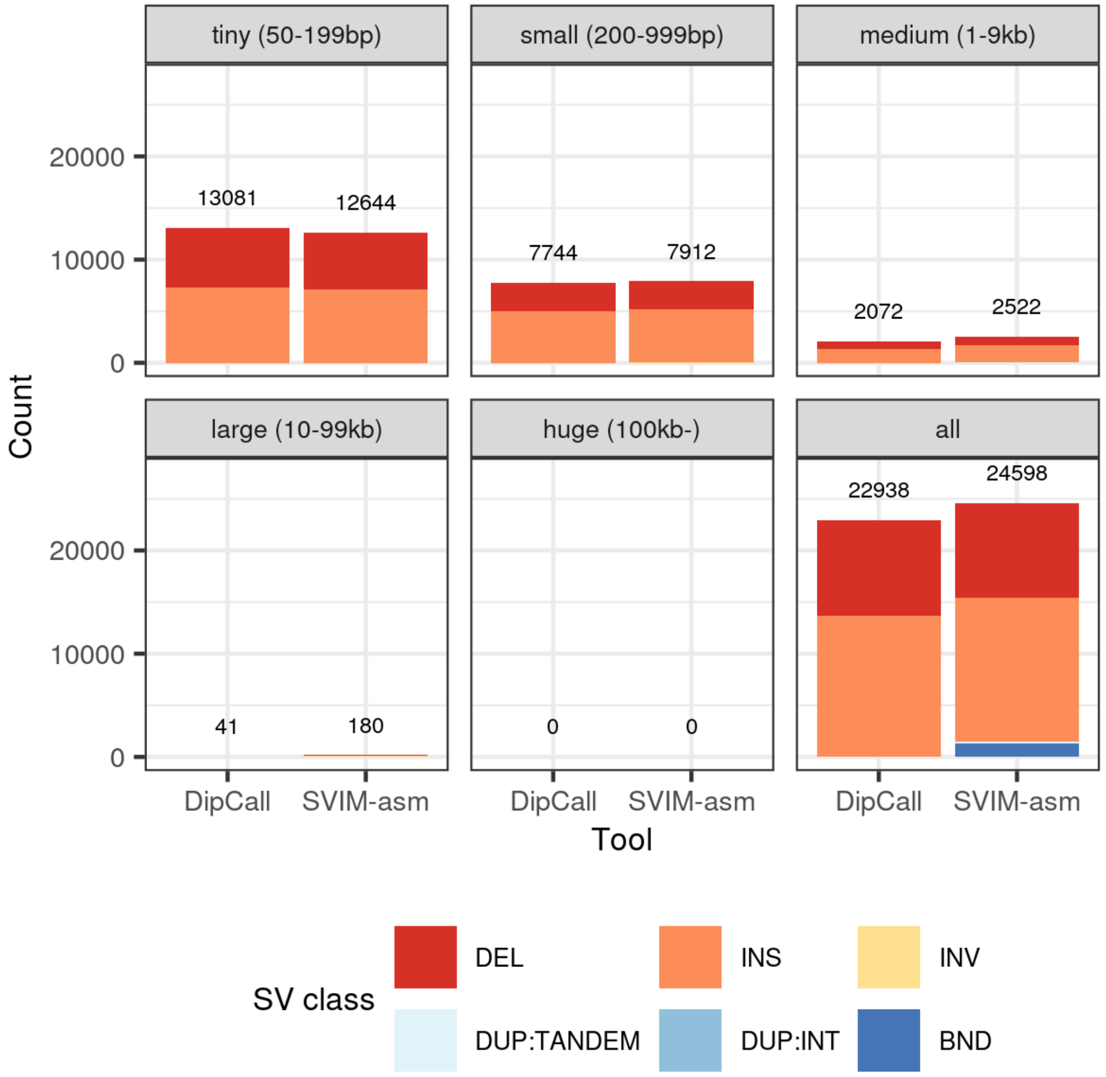


**Fig. S6 – Number of SV calls from Assembly B grouped into five size classes:** Shown are stacked bar plots of SV classes represented by different colors. Each panel represents one size class and visualizes the number of calls in that size range called by DipCall and SVIM-asm within the chromosomal contigs (1 through 22 plus X, Y and MT). The bottom right panel shows the counts for all SV calls regardless of size. As translocation breakends (BND) do not have a size, they are included only in this bottom right panel.

### Supplementary Methods

SVIM-asm analyzes genome-genome alignments in BAM format to detect six different classes of SVs. In particular, SVIM-asm searches discordant alignments in order to extract SV signatures from them. We define SV signatures as pieces of evidence pointing to the presence of an SV between the query genome assembly and the reference assembly.

Depending on the size and type of SV as well as decisions made by the alignment algorithm, discordancies can be either observed within alignment segments (intra-alignment discordancies) or between alignment segments (inter-alignment discordancies) (see Fig. S7). Intra-alignment discordancies are long alignment gaps representing deletions and insertions that can be retrieved from the CIGAR string in a BAM record. Inter-alignment discordancies, in contrast, arise when a contig is split to enable a better alignment of its segments to the reference. In this case, the termination of one alignment segment and the continuation of the alignment at another genomic position can indicate several different SV classes. Therefore, SVIM-asm classifies inter-alignment discordancies in a heuristic fashion to collect the correct type of SV signature.

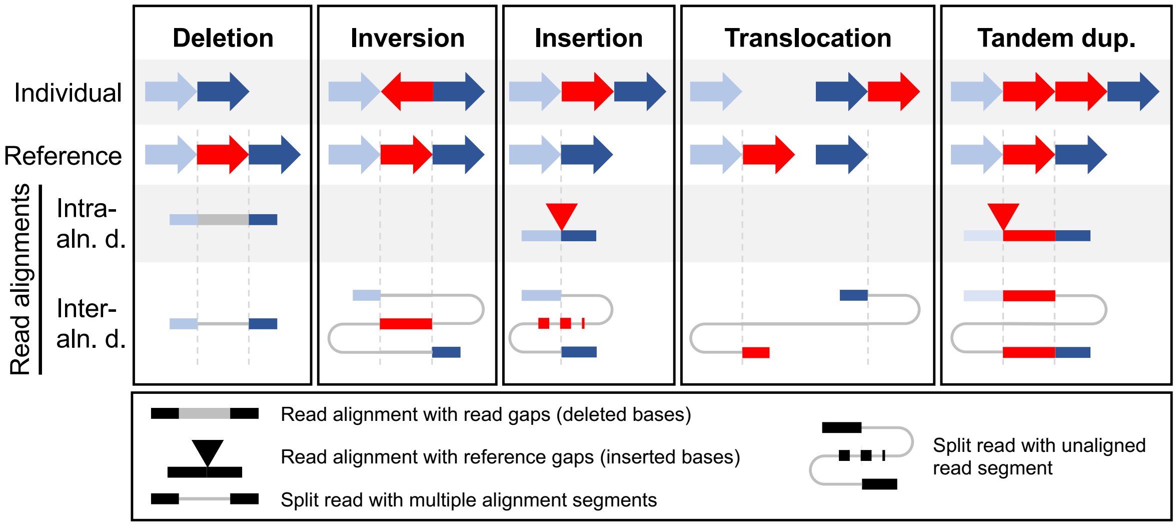


**Fig. S7 - Discordant read alignments across 5 different classes of SVs.** Structural variants between an individual genome (first row) and a reference genome (second row) lead to discordant alignments of reads from the individual genome to the reference. The discordancies can be contained either within a continuously aligned read segment (intra-alignment discordancies, third row) or between independently aligned segments of a read (inter-alignment discordancies, fourth row).
*Abbreviations: aln. - alignment, d. - discordancies, dup. - duplication*

**Diploid genome assemblies**

Diploid genome assemblies consist of two sets of contigs, one for each chromosomal haplotype. Consequently, SVIM-asm expects two input BAM alignment files as input for this type of assembly. In the first step of the pipeline (COLLECT), SV signatures are collected separately for each haplotype from individual contig alignments. While contig alignments with a low mapping quality (below 20 in our experiments) are ignored, no threshold on the number or length of contig alignments is imposed. To collect signatures from intra-alignment discordancies, the CIGAR string of each alignment segment is analyzed. Furthermore, the SA tag of primary alignments is parsed for information on supplementary alignments in order to detect inter-alignment discordancies.

In the second step of the pipeline (PAIR), signatures from opposite haplotypes are compared and paired up if sufficiently similar. To measure the similarity of two signatures $x$ and $y$, we use the edit distance (Levenshtein distance) *E(hap(R,x), hap(R,y))* between the haplotype sequences of $x$ and $y$. The haplotype sequence *hap(R,S)* of an SV signature $S$ and a reference genome sequence $R$ is the nucleotide sequence formed by applying the genomic rearrangement defined by $S$ to $R$. The edit distance measures the minimum number of operations (deletion, insertion or substitution of one character) required to transform one haplotype sequence into the other and is computed using the library edlib. To prevent that signatures from the same haplotype are matched, we enforce a very large distance instead of the actual edit distance between signatures from the same haplotype. Based on the computed distances between their haplotype sequences, the signatures are clustered using an hierarchical agglomerative clustering approach. With a low distance threshold for cutting the dendrogram, we ensure that only very similar signatures (i.e. signatures with similar haplotype sequences) from different haplotypes are clustered together.

In the third step of the pipeline (GENOTYPE), paired signatures from the two opposite haplotypes are merged into homozygous SV candidates while variants without a partner on the other haplotype are called as heterozygous SV candidates. Finally, the genotyped SVs are written out in VCF as members of one of six SV classes (OUTPUT).

**Haploid genome assemblies**

In contrast to their diploid counterparts, haploid assemblies are comprised of only a single set of contigs. For diploid organisms, this set often represents a mixture of the two haplotypes. Due to the missing second haplotype, it is not possible to estimate genotypes from haploid genome assemblies which simplifies the pipeline considerably. After the same first step (COLLECT) is applied to the assembly alignments, the PAIR and GENOTYPE steps are skipped for haploid assemblies and the detected SV signatures can be written out immediately (OUTPUT).

**Evaluation datasets**

| | ***Assembly A*** | ***Assembly B*** |
|---|---|---|
| *Authors* | Wenger et al. | Garg et al. |
| *Assembler* | Canu v1.7.1 | DipAsm |
| *Input data for contig assembly* | 29.7x PacBio CCS data (trio-binned) | 29.7x PacBio CCS data |
| *Input data for scaffolding* | No scaffolding | 28.5x Hi-C data |
| *Polishing* | Arrow v2.2.2 | No polishing |
| *Download* | Maternal: https://downloads.pacbcloud.com/public/publications/2019-HG002-CCS/asm/HG002_canu_maternal.fasta<br>Paternal: https://downloads.pacbcloud.com/public/publications/2019-HG002-CCS/asm/HG002_canu_paternal.fasta | Haplotype 1: ftp://ftp.dfci.harvard.edu/pub/hli/whdenovo/asm/NA24385-denovo-H1.fa.gz<br>Haplotype 2: ftp://ftp.dfci.harvard.edu/pub/hli/whdenovo/asm/NA24385-denovo-H2.fa.gz |
| *Citation* | Wenger, A.M., Peluso, P., Rowell, W.J. et al. Accurate circular consensus long-read sequencing improves variant detection and assembly of a human genome. Nat Biotechnol 37, 1155–1162 (2019). https://doi.org/10.1038/s41587-019-0217-9 | Garg, S., Fungtammasan, A. A., Carroll, A. et al. Efficient chromosome-scale haplotype-resolved assembly of human genomes. bioRxiv,(2019). https://doi.org/10.1101/810341 |

**SV Calling Commands**

The full SV calling and evaluation pipeline has been published at https://github.com/eldariont/assembly_sv_calling_evaluation. The commands used for calling SVs with DipCall and SVIM were as follows:

```
#DipCall
bin/dipcall.kit/run-dipcall -t 10 -x {par} {prefix} {reference} {hap1}
{hap2} > HG002.mak
make -j 40 -f HG002.mak

#SVIM
svim-asm diploid {working_dir} {bam1} {bam2} {reference} --min_sv_size
20 --tandem_duplications_as_insertions --
interspersed_duplications_as_insertions --reference_gap_tolerance 1000
--reference_overlap_tolerance 1000 --query_gap_tolerance 2000 --
query_overlap_tolerance 2000 --max_edit_distance 200 --sample HG002 --
query_names
```

**Evaluation Commands**

The full SV calling and evaluation pipeline has been published at https://github.com/eldariont/assembly_sv_calling_evaluation. The commands used for comparing SV calls with the benchmark set were as follows:

```
#compare calls with benchmark set
truvari bench -f {reference} -b {truth} --includebed {truth_regions} -o
{out_dir} --giabreport --passonly -r 1000 -p 0 -c {calls}

#compare genotypes of true positive calls with benchmark set
python workflow/scripts/compare_genotypes.py {tp_call} {tp_base} {fp}
{fn} > {results}
```

The compare_genotypes.py script can be accessed here:
https://github.com/eldariont/assembly_sv_calling_evaluation/blob/master/workflow/scripts/compare_genotypes.py

**Data availability**

The SV calls by SVIM and DipCall and the SV benchmark set are available at https://owww.molgen.mpg.de/~svim/asm/svim-asm-data.zip. As reference genome, hs37d5 was used which can be downloaded here:
ftp://ftp-trace.ncbi.nih.gov/1000genomes/ftp/technical/reference/phase2_reference_assembly_sequence/hs37d5.fa.gz
